# Guild-level microbiome signature associated with COVID-19 severity and prognosis

**DOI:** 10.1101/2022.09.18.508418

**Authors:** Mingquan Guo, Guojun Wu, Yun Tan, Yan Li, Xin Jin, Weiqiang Qi, XiaoKui Guo, Chenhong Zhang, Zhaoqin Zhu, Liping Zhao

## Abstract

COVID-19 severity has been associated with alterations of the gut microbiota. However, the relationship between gut microbiome alterations and COVID-19 prognosis remains elusive. Here, we performed a genome-resolved metagenomic analysis on fecal samples collected from 300 in-hospital COVID-19 patients at time of admission. Among the 2,568 high quality metagenome-assembled genomes (HQMAGs), Redundancy Analysis identified 33 HQMAGs which showed differential distribution among mild, moderate, and severe/critical severity groups. Random Forest model based on these 33 HQMAGs classified patients from different severity groups (average AUC = 0.79). Co-abundance network analysis found that the 33 HQMAGs were organized as two competing guilds. Guild 1 harbored more genes for short-chain fatty acid biosynthesis, and fewer genes for virulence and antibiotic resistance, compared with Guild 2. Random Forest regression showed that these 33 HQMAGs at admission had the capacity to predict 8 clinical parameters, which are predictors for COVID-19 prognosis, at Day 7 in hospital. Moreover, the dominance of Guild 1 over Guild 2 at admission predicted the death/discharge outcome of the critical patients (AUC = 0.92). Random Forest models based on these 33 HQMAGs classified patients with different COVID-19 symptom severity, and differentiated COVID-19 patients from healthy subjects, non-COVID-19, and pneumonia controls in three independent datasets. Thus, this genome-based guild-level signature may facilitate early identification of hospitalized COVID-19 patients with high risk of more severe outcomes at time of admission.

## Introduction

Coronavirus disease 2019 (COVID-19), caused by novel severe acute respiratory syndrome coronavirus 2 (SARS-CoV-2), has been a worldwide pandemic with heavy toll to human health and economy. Over 576 million people have been infected by SARS-CoV-2, with over 6 million deaths globally ^1^. Angiotensin-converting enzyme 2 (ACE-2), which is distributed in multiple tissues and widely expressed on the luminal surface of the gut, has been identified as a vital entry receptor of SARS-CoV-2 for promoting viral infection and replication^2^. This can impair gut barrier and induce inflammation, which may disrupt the gut microbiome, contributing to cytokine storm and sepsis in already compromised patients with COVID-19^2^.

Recent studies have showed that dysbiosis of the gut microbiome and its related metabolites are closely associated with COVID-19 diseases. These studies reveal the overall difference in the gut microbial composition between COVID-19 patients and healthy controls^3–11^, association of microbial taxa and metagenomic functions with disease severity^3,8,9,11^ and persistent dysbiosis of the gut microbiota after recovery^3^. The enrichment of pathobionts and depletion of beneficial microbes have been reported to be related to the disease severity in COVID-19^4,7^. However, these studies have suffered from small sample size, lack of cross study validation and missing microbiome signature at admission for prognosis of COVID-19 in hospitalized patients^4,8–11^. In addition, the reported findings are constrained by analyzing the microbiome at low resolution levels, such as species, genus or even phylum or broad metagenomic functional categories^3–11^. In the gut microbial ecosystem, the strains/genomes are the minimum responding units to environmental perturbations and their response and contributions to the host are not constrained by taxonomy, even in the same species^12^.

In this study, we obtained high-quality metagenome-assembled genomes (HQMAGs) from metagenomically sequenced fecal samples collected from 300 in-hospital COVID-19 patients with mild, moderate, severe, and critical severities at time of admission. We identified a guild-level microbiome signature of 33 HQMAGs. This signature classified patients with different severity, predicted clinical parameters related with prognosis after 1 week in hospital, and the death/discharge outcome of critical patients. The capacity of this signature for classifying COVID-19 patients with different level of severity and differentiating COVID-19 patients from healthy individuals, non-COVID-19 and pneumonia control was validated in three independent datasets.

## Results

### Overall structural changes of the gut microbiome were associated with disease severity in COVID-19 patients at admission

From May to September 2020, we collected 330 stool samples from 300 in-hospital patients with COVID-19 confirmed by positive SARS-CoV2-2 RT-qPCR result. Among the 330 samples, 297 were collected from 297 patients at admission, and 33 samples were collected from 29 patients during their hospitalization (Table S1). To profile the gut microbiome, metagenomic sequencing was performed on all the 330 stools samples. To achieve strain/subspecies level resolution, we reconstructed 2,568 non-redundant HQMAGs (two HQMAGs were collapsed into one if the average nucleotide identity (ANI) between them was > 99%) from the metagenomic dataset. The HQMAGs accounted for more than 77.17% ± 0.23% (mean ± S.EM.) of the total reads and were used as the basic variables in the further microbiome analysis.

The 296 patients with metagenomic dataset at admission (one sample was discarded due to low mapping rate of the reads against HQMAGs) were classified into the mild (N=88), moderate (N=196), severe (N=5) and critical (N=7) groups based on their symptoms. Due to the limited sample size of severe and critical patients, we combined these two groups as one group in the following analysis. There were significant differences in age between the patients with mild, moderate, and severe/critical symptoms (Kruskal-Wallis test, *P* = 1.6×10^-14^) i.e., the more severe symptoms the patients had, the older they were (Fig. S1). There was no difference in gender among the 3 groups (chiq-square test, *P* = 0.22).

At admission, in the context of beta-diversity based on the Bray-Curtis distance, Principal Coordinate Analysis (PCoA) revealed separations of the gut microbiota along PC1, which was in accord with the severity of symptoms (Fig. 1A-C). A marginal PERMANOVA test, including disease severity and age, showed that disease severity was significantly associated with the overall gut microbial composition (R^2^ = 0.012, *P* = 0.0002,) but age was not (R^2^ = 0.0032, *P* = 0.50). This showed that in our dataset when disease severity was held constant, the age effect on gut microbiome was not significant. The pairwise comparisons between the 3 different severity groups via PERMANOVA test showed that gut microbial composition of the patients was significantly different from each other (mild vs. moderate: R^2^ = 0.0081, *P* = 0.0001; mild vs. severe/critical: R^2^ = 0.026, *P* = 0.0001; moderate vs. severe/critical: R^2^ = 0.0079, *P* = 0.0099). The distance between the mild and moderate groups was significantly smaller than that between the mild and the severe/critical groups (Fig. S2), which showed that the gut microbiota of severe/critical group was more different from the mild group as compared with the moderate group. Regarding to alpha-diversity, Shannon index was highest in the mild group, followed by the moderate group and lowest in the severe/critical group (Fig. 1D, mild vs. moderate: *P* = 0.0046, mild vs severe/critical: *P* = 0.0046, moderate vs. severe/critical: *P* = 0.086), which showed a continuous reduction in gut microbial diversity with the increase of symptom severity. These results showed that the overall gut microbial structure was associated with the severity of symptoms of COVID-19 patients.

**Figure 1.**
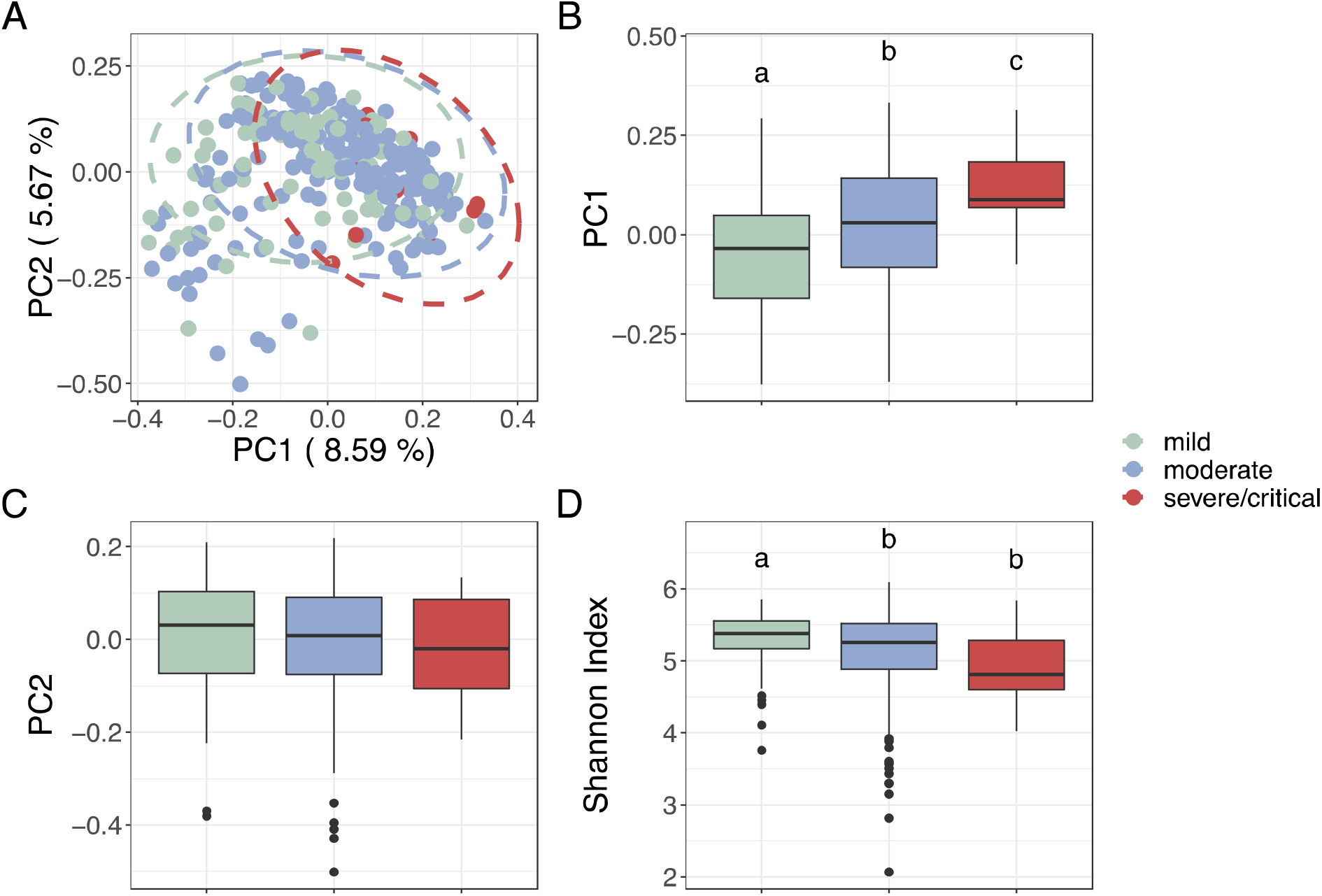
The overall structural variations of gut microbiota at admission associated with disease severity in hospitalized COVID-19 patients. (A) Principal Coordinate Analysis based on Bray-Curtis calculated from abundance of the 2,568 genomes. (B) and (C) Comparison of the PC1 and PC2. (D) Comparison of alpha-diversity as indicated by Shannon index. Data points not sharing common compact letters were significantly different from each other (p < 0.05). Boxes show the medians and the interquartile ranges (IQRs), the whiskers denote the lowest and highest values that were within 1.5 times the IQR from the first and third quartiles, and outliers are shown as individual points. Kruskal-Wallis test followed by Dunn’s post hoc (two-sided) was applied to compare the groups. Compact letters reflect the significance of the test (P < 0.05). Mild: n = 88; moderate n = 196, severe/critical n = 12.

### Two competing guilds were associated with disease severity of hospitalized COVID-19 patients at admission

Specific HQMAGs that were associated with the COVID-19 symptom severity were identified by redundancy analysis (RDA) (Fig. S3). Out of the 2,568 HQMAGs, we found that 48 HQMAGs had at least 5% of their variability explained by the constraining variable, i.e., the three severity groups. Among the 48 HQMAGs, 17 were significant higher in the mild group as compared with the moderate and severe/critical groups and they showed a continuous decrease along the symptom severity (Fig. 2A). These 17 HQMAGs included 5 from *Faecalibacterium prausnitzii,* 3 from *Romboutsia timonensis, 2* each from *Ruminococcus, Clostridium* and 1 each from Acutalibacteraceae, *Allisonella histaminiformans, Coprococcus,* Lachnospiraceae and *Negativibacillus.* The abundance of 31 out of the 48 RDA identified HQMAGs were higher in the severe/critical group as compared with the mild and the moderate groups. Among these 31 HQMAGs, 16 showed significant difference between the three groups. These 16 HQMAGs included 4 from *Enterococcus, 2* from *Lactobacillus,* 1 each from Acutalibacteraceae, *Akkermansia muciniphila, Anaerotignum, Barnesiella intestinihominis, Clostridium_M bolteae, Dore, Intestinibacter bartlettii,* Lachnospiraceae *,Phascolarctobacterium faeciu* and *Ruthenibacterium lactatiformans. We* then focused on the 33 HQMAGs which were both identified by RDA analysis and were significantly different between the 3 severity groups. Next, we developed a machine learning classifier based on Random Forest (RF) algorithm to see if we could classify patients from different severity groups based on the 33 HQMAGs. Receiver operating characteristic curve analysis showed a power with area under curve (AUC) of 0.76, 0.84 and 0.76 to classify mild vs. moderate, mild vs. sever/critical, and moderate vs. severe/critical (Fig. 2B). These results showed that these 33 HQMAGs had the capacity to differentiate the different symptom severity groups of COVID-19 patients.

**Figure 2.**
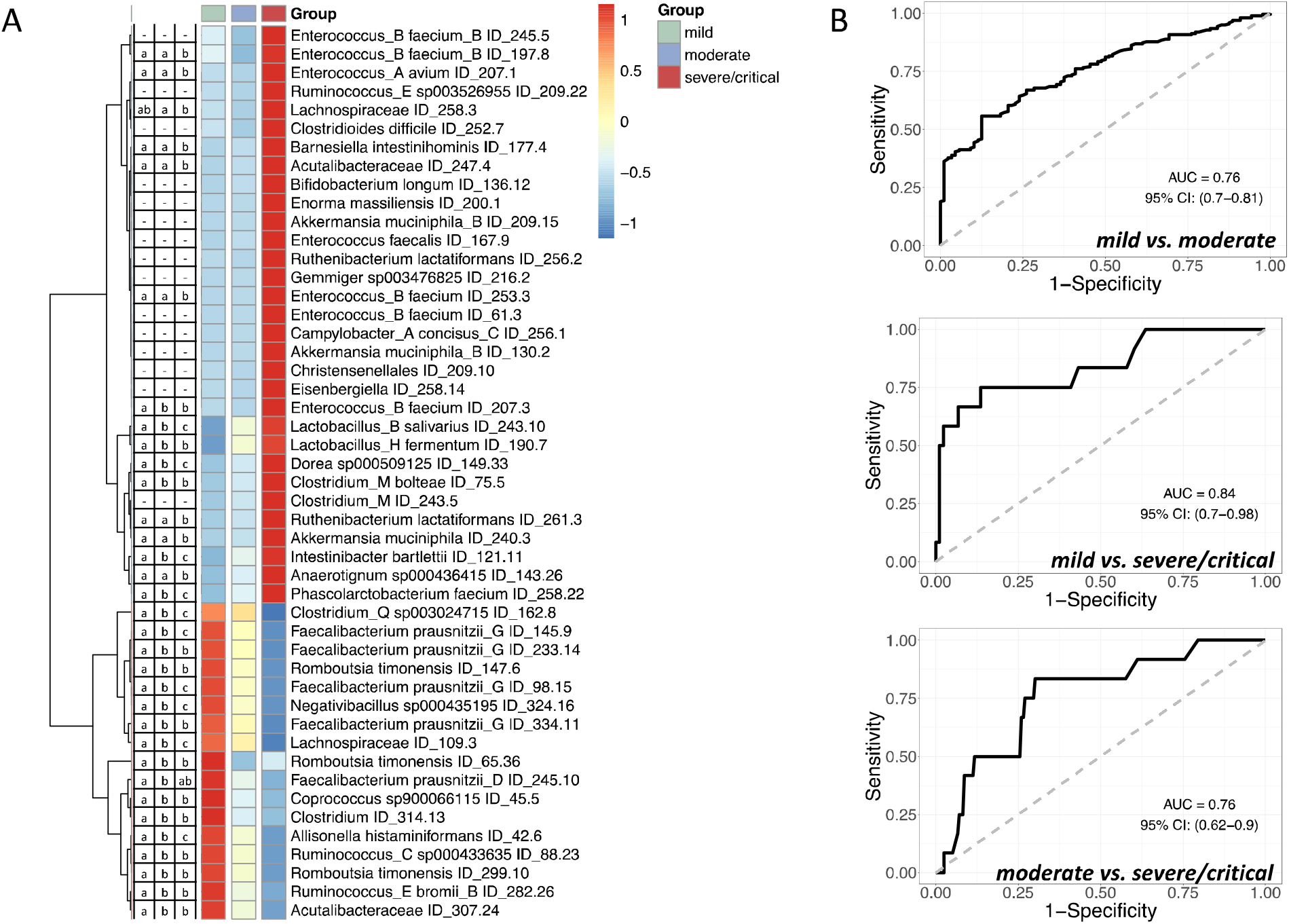
Differentially abundant gut microbial genomes classify patients with different COVID-19 symptom severity. (A) The heatmap of 48 high quality metagenome-assembled genomes (HQMAGs) identified by redundancy analysis (RDA). RDA analysis was conducted based on the Hellinger transformed abundance of all the HQMAGs and use the three symptom severity groups as environmental variables. HQMAGs with at least 5% of the variability in their abundance explained by constrained axes were selected. The heatmap shows the mean abundance of each HQMAGs in each group. The abundance was scaled across each row. The HQMAGs were clustered based on Euclidean distance and complete linkage. Kruskal-Wallis test followed by Dunn test (two-sided) was used to test the differences between the 3 severity groups. Compact letters reflect the significance as P < 0.05. (B) The area under the ROC curve (AUC) of the Random Forest classifier based on the 33 HQMAGs to classify different COVID-19 symptom severity. Leave-one-out cross validation was applied.

As bacteria in the gut ecosystem are not independent but rather form coherent functional groups (a.k.a “guilds”) to interact with each other and affect host health^14^, we applied co-abundance analysis on these 33 HQMAGs to explore the interactions between them and to find potential guilds structure. Interestingly, the 33 HQMAGs organized themselves into two guilds—the 17 HQMAGs with significantly higher abundance in mild group were positively interconnected with each other and formed as Guild 1. The 16 severe/critical group enriched HQMAGs were positively correlated with each other as Guild 2 (Fig 3A). Meanwhile, there were only negative correlations between the two guilds, suggesting a potentially competitive relationship between the two guilds.

**Figure3.**
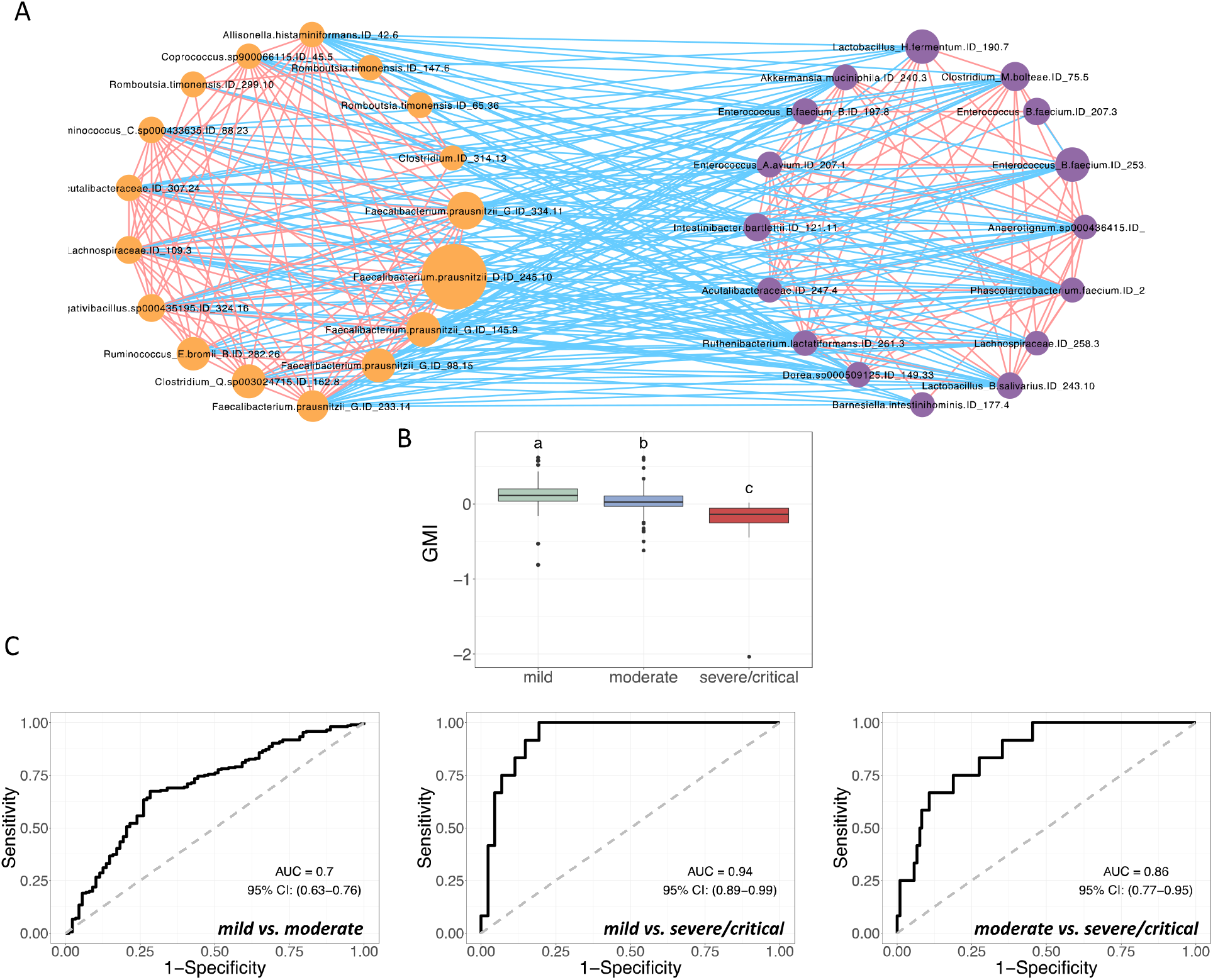
Two competing guilds associated with the symptom severity of COVID-19 patients. **(A)** Co-abundance network of the HQMAGs reflects two competing guilds. The co-abundance correlation between the HQMAGs were calculated using Fastspar, n= 296 subject. All significant correlations with BH-adjusted p < 0.05 were included. Edges between nodes represent correlations. Red and blue colors indicate positive and negative correlations, respectively. Node size indicates the average abundance of the HQMAGs in 296 samples. Node color indicates Guild 1 (orange) and Guild 2 (purple), respectively. (B) Comparison of guild-level Microbiome Index (GMI). Data points not sharing common compact letters were significantly different from each other (p < 0.05). Boxes show the medians and the interquartile ranges (IQRs), the whiskers denote the lowest and highest values that were within 1.5 times the IQR from the first and third quartiles, and outliers are shown as individual points. Kruskal-Wallis test followed by Dunn’s post hoc (two-sided) was applied to compare the groups. Compact letters reflect the significance of the test (P < 0.05). Mild: n = 88; moderate n = 196, severe/critical n = 12. (C) The Guild-level Microbiome Index (GMI) supports classification for different COVID-19 symptom severity. Leave-one-out cross validation was applied for each model.

To explore the genetic basis underling the associations between the two guilds and the symptom severities, we performed genome-centric analysis of the metagenomes of the two competing guilds. A previous study showed that the lack of short chain fatty acids (SCFAs) significantly correlated with disease severity in COVID-19 patients^15^. Regarding to the terminal genes for the butyrate biosynthetic pathways (i.e., *but, buk, atoA/D* and *4Hbt*)^16^, 7 HQMAGs in Guild 1 harbored but genes while only 1 HQMAGs in Guild 2 had this gene (Fisher’s exact test *P* = 0.039) (Fig. S4). Four HQMAGs in Guild 1 harbored *buk* genes while no HQMAGs in Guild 2 had this gene (Fisher’s exact test *P* = 0.10). The other butyrate biosynthetic terminal genes were not found in the HQMAGs in both guilds. The numbers of HQMAGs encoding genes for acetate and propionate production were similar in the 2 guilds (Fig. S4). From the perspective of pathogenicity, although in both guilds there were 12 HQMAGs encoding virulence factor (VF) genes, the number of VF genes (17 in Guild 1, 58 in Guild2) and the VF categories (3 in Guild 1, 5 in Guild 2) were higher in Guild 2 as compared with Guild 1 (Fig. S5A). In terms of antibiotic resistance genes (ARGs), 3 genomes in Guild 1 encoded 10 ARGs and 5 genomes in Guild 2 encoded 14 ARGs (Fig. S5B). Taken together, these data showed that the two competing guilds had different genetic capacity with Guild 1 being more beneficial and Guild 2 more detrimental. Thus, the genetic difference between the two guilds may help explain their associations with the disease severity of COVID-19 patients.

We then calculated the Guild-level microbiome index (GMI) based on the average abundance difference between Guild 1 and Guild 2 to reflect the dominance of Guild 1 over Guild 2. At admission, the GMI was highest in the mild group, followed by the moderate group and was lowest in the severe/critical group (Fig. 3B, mild vs. moderate: *P* = 2.46 × 10^-7^, mild vs severe/critical: *P* = 6.57 × 10^-9^, moderate vs. severe/critical: *P* = 1.59 × 10^-4^). The GMI reached an AUC of 0.7 to differentiate mild and moderate groups, an AUC of 0.94 to differentiate mild and sever/critical groups and an AUC of 0.86 to differentiate moderate and severe/critical groups (Fig. 3C). This result indicates the feasibility to use the GMI as a biomarker to differentiate the different symptom severity groups of COVID-19 patients.

### Gut microbiome signature was associated with the prognosis of COVID-19 in hospitalized patients

To explore whether our microbiome signature at admission is associated with the prognosis of COVID-19 patients during hospitalization, we applied RF algorithm to regress the abundance of the 33 HQMAGs at admission with 72 different clinical parameters at Day 7 in hospital. Nine clinical parameters at day 7 were significantly associated with the abundances of the 33 HQMAGs at admission as evidenced by significantly positive correlations between the measured values and the predicted values from the regression model (Pearson correlation: BH adjusted *P* < 0.1 and R > 0) (Fig 4A). In addition, 9 and 11 clinical parameters at Day 7 showed significantly (Spearman correlation: BH adjusted *P* < 0.1) positive and negative correlations with the GMI values at admission, respectively (Fig. 4A). Regarding immune indicators, interleukin (IL)-5 is secreted chiefly by Th2 cells and essentially anti-inflammatory but also involved in several allergic responses^17^. Some studies have revealed a higher level of IL-5 in severe cases compared with mild cases^18,19^. However, some studies have showed that IL-5 have no correlations with COVID-19 and showed no differences between different severity groups^20,21^. Like IL-5, the association between IL-12p70 and COVID-17 remain elusive as some studies have revealed elevated IL-12p70 in COVID-19 but others not^22–24^. Here, we found positive correlations between the GMI at admission and these two ILs after 1 week. The effects of microbiome on particular cytokines and the subsequent influences on COVID-19 need further studies. Coagulation disorder occurred at the early stage of COVID-19 infection^25^. D-Dimer, Fibrinogen (Fg) and fibrin degradation product (FDP) increased in COVID-19 patients and were correlated with clinical classification^25,26^. Moreover, elevated D-Dimer, Fg and FDP are significant indictors of severe COVID-19 and poor prognosis^25–28^. Here, a higher GMI at admission was correlated with lower D-Dimer, Fg and FDP after 1 week. Regarding hemogram indicators, the degree of lymphopenia is an effective and reliable indicator of the severity and hospitalization in COVID-19 patients^29,30^. Both blood lymphocyte percentage (LYMPH%) and absolute count (LYMPH #) are strong prognostic biomarkers in hospitalized patients with severe COVID-19^31^. Blood LYMPH% has been reported to be inversely associated with severity and poor prognosis in COVID-19^30^. In contrast to LYMPH%, neutrophils percentage (Neu %) were higher in COVID-19 patients with severer symptoms and directly associated with poor prognosis^31,32^. Here, the GMI at admission was positively correlated with LYMPH % and LYMPH #, and negatively correlated with Neu % after 1 week. Regarding biochemical indicators, compared with health subjects, total cholesterol (TC) was significant lower in COVID-19 patients and decreased with increasing severity^33,34^. A Meta-analysis showed that the reduction of TC was significantly associated with the increased mortality in COVID-19 patients and TC might assist with early risk stratification^34^. Hypocalcemia has been reported to be common in COVID-19 patients^35^. Total calcium (CA) at admission showed an inversely relationship with the mortality rate in COVID-19 patients^35^. Among the abnormal liver biochemical indictors at admission, abnormal albumin (ALB) was the most common^36^. A declined ALB level was associated with the disease severity of COVID-19^36^. Higher total bilirubin (TBIL) were associated with a significant increase in the severity of COVID-19 infection^37^. Moreover, COVID-19 patients with an elevated TBIL at admission had a higher mortality^37^. In addition to TBIL, increased direct bilirubin (DBIL) has been reported as an independent indicator for complication and mortality in COVID-19 patients^38^. Particularly, DBIL levels at Day 7 of hospitalization are advantageous in predicting the prognosis of COVID-19 in severe/critical patients^38^. Lactate dehydrogenase (LDH) have been associated with worse outcomes in viral infection. A meta-analysis showed that LDH could be used as a COVID-19 severity marker and a predictor of survival^39^. In COVID-19, elevation in glucose level has been linked with major steps of the life cycle of the virus, progression of the disease, and presentation of symptoms^40^. It has been reported that during hospitalization patients with well-controlled glucose had markedly lower mortality compared to individuals with poorly controlled glucose^41^. Here, the GMI at admission was positively correlated with TC, CA and ABL, and negatively correlated with TBIL, DBIL, LDH and glucose after 1 week. These results suggest that gut microbiome signature in early stage may reflect the clinical outcomes of COVID-19 in hospitalized patients.

**Figure4.**
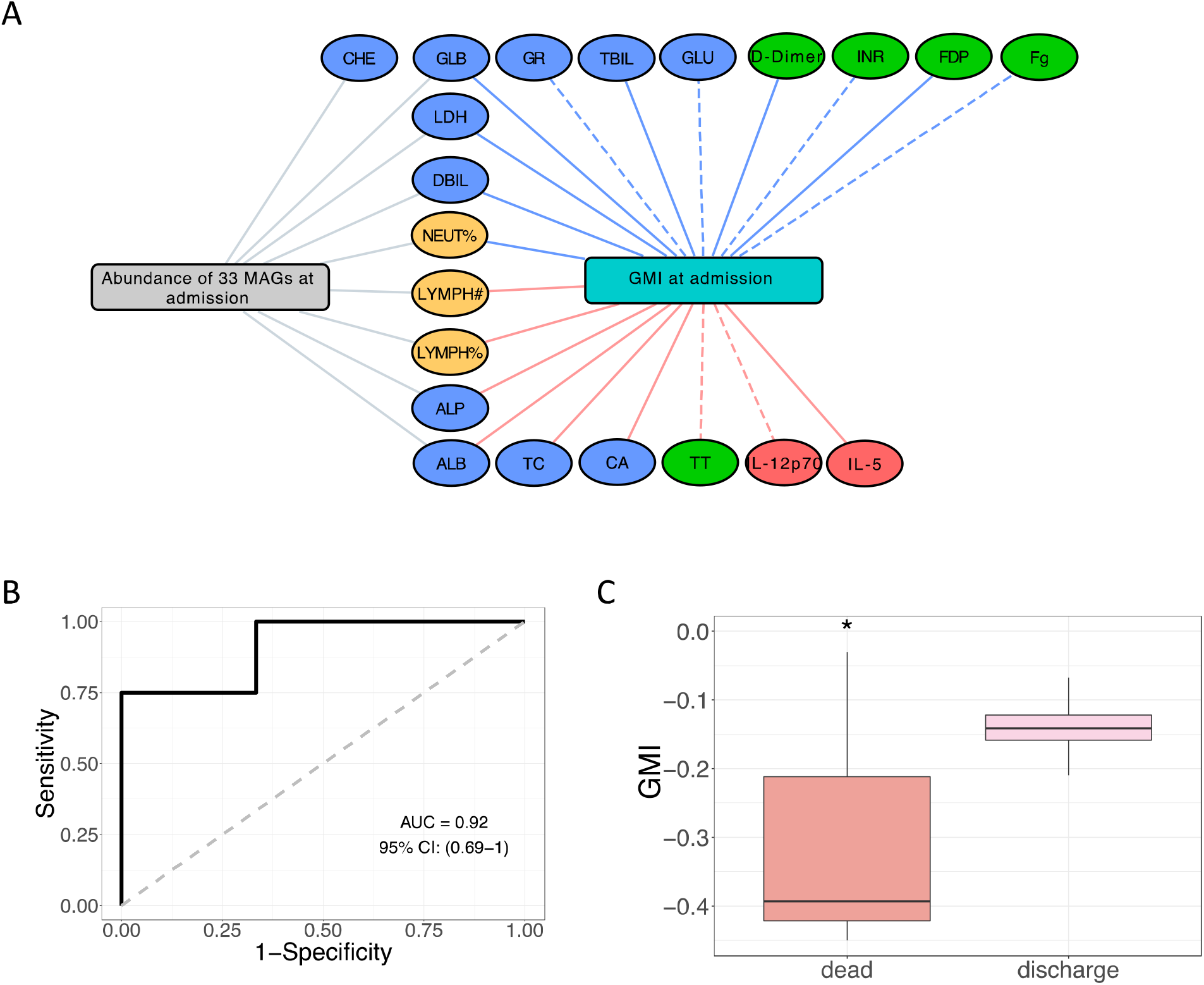
The two competing guilds at admission associated with severity of COVID-19 in hospitalized patients at Day 7 after admission and outcome in critical patients. (A) The network shows the correlations between the microbiome signature at admission and clinical parameters of COVID-19 in hospitalized patients at Day 7. Random Forest (RF) regression with leave-one-out cross-validation was used to explore the associations between the 33 genomes at admission and the clinical parameters at Day 7. Edges in gray show the clinical parameters with significantly positive Pearson’s correlations between the measured values and the predicted value from the RF model. Spearman correlation was used to explore the associations between the Guild-level Microbiome Index (GMI) at admission and the clinical parameters at day 7. Edges in red and blue show the significantly positive and negative correlations, respectively. The color of the nodes: blue: biochemical indicators, green: coagulation indicators; red: immune indicators; yellow: hemogram indicators. Line type: solid: BH adjusted P < 0.05, dashed: BH adjusted P < 0.1. CHE: serum cholinesterase, GLB: globulin, LDH: lactate dehydrogenase, DBIL: direct bilirubin, NEUT%: neutrophils percentage, LYMPH #: lymphocyte count, LYMPH %: lymphocyte percentage, ALP: Alkaline phosphatase, ALB: albumin; GR: glutathione reductase; TC: total cholesterol, TBIL: total bilirubin, CA: total calcium, Glu: glucose, TT: thrombin time; IL-12p70: Interleukin −12p70, FDP: fibrin degradation products, Fg: fibrinogen(B) The area under the ROC curve (AUC) of the Random Forest classifier based on the 33 HQMAGs at admission to classify the outcome (death/discharge) of critical COVID-19 in hospitalized patients. Leave-one-out cross validation was applied. (C) Guild-level Microbiome Index (GMI) at admission associate with the ends of critical COVID-19 patients. Two-sided Mann-Whitney test was applied. Death n =3, discharge n = 4. * P < 0.05.

Moreover, in our cohort, 3 patients were dead, and they were all in the critical group at admission. Compared with the other 4 discharged critical patients, the 3 dead patients were significantly younger (Fig. S6). To explore if the microbiome signature at admission can predict the outcome of the critical COVID-19 patients, we applied RF algorithm to classify death/discharge with the abundance of the 33 HQMAGs at admission, and the model reached an AUC of 0.92 (Fig. 4B). In addition, we found that the GMI values at admission of the 3 dead patients were significantly lower than those of the 4 discharged critical patients (Fig. 4C). This suggests an association between the microbiome signature with the final outcome in critical hospitalized COVID-19 patients. Although interesting, this result should be interpreted with caution given the small sample size.

### The microbiome signature was validated in independent studies

We then asked that whether this genome-based microbiome signature would be applicable in other COVID-19 cohorts. To answer this question, we used the genomes of the 33 HQMAGs as reference genomes to perform read recruitment analysis, which is a commonly used way to estimate abundances of reference genomes from metagenomes^42,43^. In an independent study, which included 24 mild/moderate and 14 severe/critical COVID-19 patients^9^, we validate the associations between the microbiome signature and the different COVID-19 severity. In this validation dataset, the two groups of patients had even distributions of age, gender, and comorbidities, which avoid potential biases for our validation. On average, the 33 HQMAGs accounted for 4.39 % ± 0.90% (mean ± S.E.M) of the total abundance of the gut microbial community. In the context of beta-diversity measured via the Bray-Curtis distance, the composition of the microbiome signature between the mild/moderate and severe/critical COVID-19 patient were significantly different (Fig. 5A). Next, we developed a classifier based on RF algorithm to see if we could classify patients from different severity groups based on the 33 HQMAGs. ROC curve analysis showed a power with AUC of 0.89 to classify mild/moderate and severe/critical groups (Fig. 5B). In addition, regrading GMI and the abundance of the 2 guilds, we found that GMI and the abundance of Guild 1 were significantly higher in the mild/moderate patients, while the abundance of Guild 2 was significantly higher in the severe/critical patients (Fig. 5C). These results validated our findings of the associations between the genome-resolved microbiome signature and the COVID-19 disease severity.

**Figure 5.**
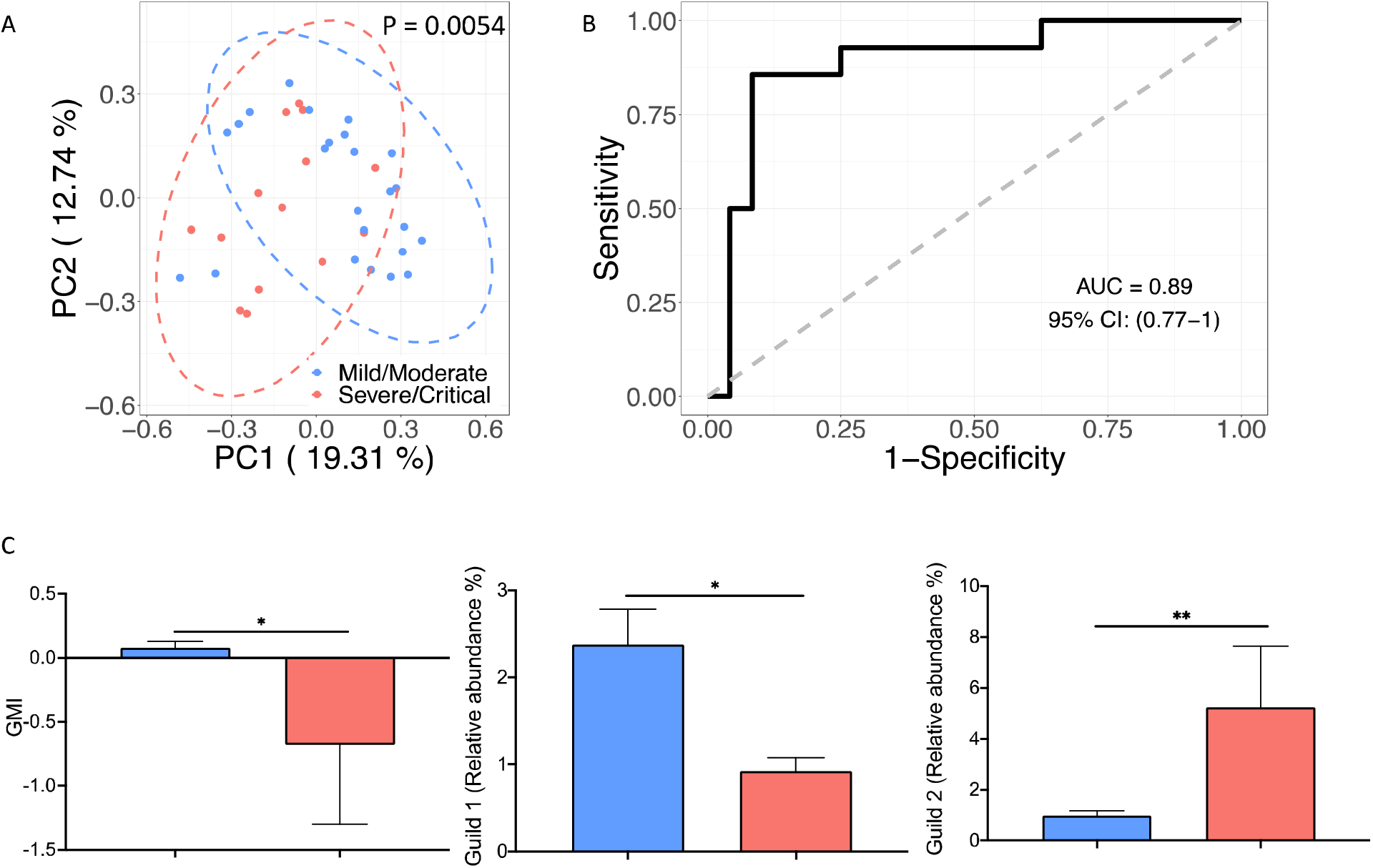
The genome-based microbiome signature enables to classify COVID-19 patient from different severity groups in an independent dataset. (A) Principal Coordinate Analysis based on Bray-Curtis distance calculated from the abundance of the 33 HQMAGs. PERMANOVA test showed significant differences in the composition of the 33 HQMAGs between the two groups(B) The area under the ROC curve (AUC) of the Random Forest classifier based on the 33 HQMAGs to classify mild/moderate and severe/critical COVID-19 patients. Leave-one-out cross validation was applied. (C) Significant differences in Guild-level Microbiome index (GMI) and abundances of Guild 1 and Guild 2 between mild/moderate and severe/critical COVID-19 patients. The barplot summarized the mean and S.E.M. Mann-Whitney test (two-sided) was applied to compare the groups. Mild/Moderate n = 24, Severe/Critical n =14. * P < 0.05, ** P < 0.01

Since this microbiome signature was associated with COVID-19 disease and was able to classify COVID-19 severity, we are interested to find out if it would be able to classify COVID-19 and non-COVID-19 controls as well. We included metagenomic sequencing data from 66 COVID-19 patients (first sample after admission), 69 age- and sex-matched non-COVID-19 controls and 9 community-acquired pneumonia controls but were negative for COVID-19 from the study conducted by Zhang et al.^7^. The genomes of the 33 HQMAGs were used as reference genomes to perform read recruitment analysis. On average, the 33 HQMAGs accounted for 3.47% ± 0.40% (mean ± S.E.M) of the total abundance of the gut microbial community. Based on the Bray-Curtis distance, the PCoA plot revealed separations between non-COVID-19 control and the other two groups (Fig. 6A and B). Though no significant separation were found between COVID-19 and pneumonia control along PC1, PERMANOVA test showed significant differences between the 3 groups pair-wisely (Fig. 6C). These results suggest that the SARS-COV-2 infection is associated with altered composition of the 33 HQMAGs. Next, we developed a classifier based on RF algorithm to see if we could classify the subjects from the 3 groups based on the abundance of the 33 HQMAGs. ROC curve analysis showed a power with AUC of 0.81, 0.8 and 0.91 to classify COVID-19 vs. non-COVID-19, COVID-19 vs. pneumonia control and non-COVID-19 vs. pneumonia control (Fig. 6D). In addition, we found that GMI were highest in non-COVID-19, followed by COVID-19 and lowest in the pneumonia control (Fig. 6E, COVID-19 vs. non-COVID-19: P = 0.11, COVID-19 vs. pneumonia control: P = 0.018, non-COVID-19 vs. pneumonia control: P = 0.0016). Non-COVID-19 had highest abundance of Guild 1 and pneumonia control had highest abundance of Guild 2 (Fig. 6E). The abundance of Guild 1 and Guild 2 in the COVID-19 group were in the middle across the 3 groups. Such differences of GMI across the 3 groups are supposed to validate our finding that the higher GMI the better healthy condition because subjects from non-COVID-19 are supposed to be the healthiest, most of the COVID-19 subjects (47 out of 66) were mild and moderate, and subjects from pneumonia control are supposed to be the least healthy. In addition to this dataset, we included metagenomic sequencing data from 46 COVID-19 patients and 19 age- and sex-matched healthy controls from the study conducted by Li et al.^44^. On average, the 33 HQMAGs accounted for 1.61 % ± 0.12% (mean ± S.E.M) of the total abundance of the gut microbial community. Based on the Bray-Curtis distance, the PCoA plot revealed a separation between COVID-19 and healthy subjects (Fig. S7A). The classifier based on RF algorithms with the abundance of the 33HQMAGs as input variables showed a power with AUC of 0.75 to classify COVID-19 and healthy controls (Fig. S7B). Compared with healthy controls, COVID-19 patients had significantly lower GMI and abundance of Guild 1 but higher abundance of Guild 2. These showed that the microbiome signature is relevant with the host health and has the capacity to be used as biomarkers to differentiate the COVID-19 subjects from healthy subjects, non-COVID-19 and pneumonia controls.

**Figure 6.**
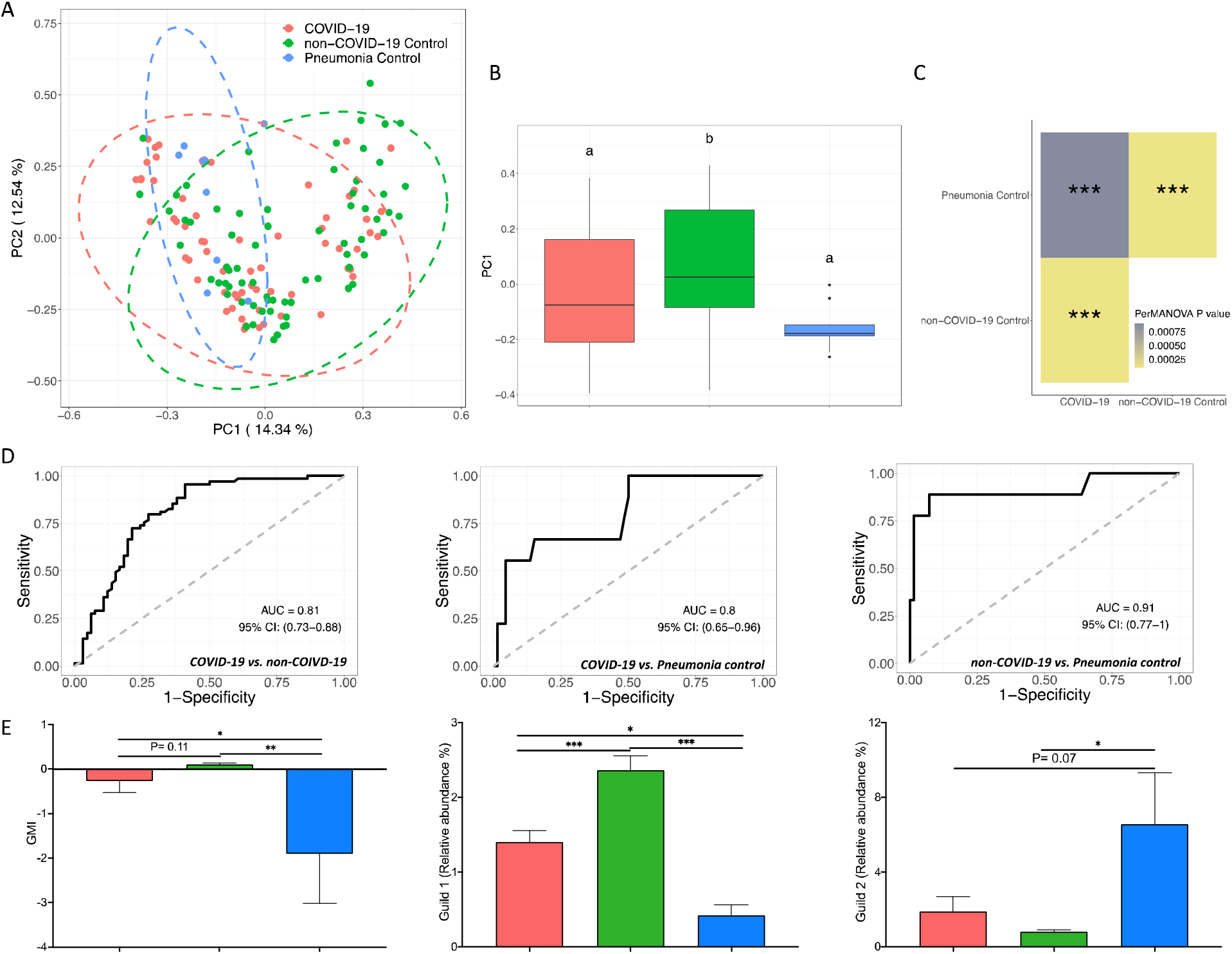
The genome-based microbiome signature enabled classification between COVID-19, non-COVID-19 and pneumonia in an independent dataset. (A) Principal Coordinate Analysis based on Bray-Curtis distance calculated from the abundance of the 33 HQMAGs. (B) Comparison of the PC1. Data points not sharing common compact letters were significantly different from each other (p < 0.05). Boxes show the medians and the interquartile ranges (IQRs), the whiskers denote the lowest and highest values that were within 1.5 times the IQR from the first and third quartiles, and outliers are shown as individual points. (C) PERMANOVA test showed significant differences in the composition of the 33 HQMAGs between COVID-19, non-COVID-19 and pneumonia subjects. *** P < 0.001 (D) The area under the ROC curve (AUC) of the Random Forest classifier based on the 33 HQMAGs to classify COVID-19, non-COVID-19 and pneumonia subjects. Leave-one-out cross validation was applied. (E) Significant differences in Guild-level Microbiome index (GMI) and abundances of Guild 1 and Guild 2 between COVID-19, non-COVID-19 and pneumonia subjects. The barplot summarized the mean and S.E.M. Kruskal-Wallis test followed by Dunn’s post hoc (two-sided) was applied to compare the groups. Compact letters reflect the significance of the test (P < 0.05). COVID-19 n = 66, non-COVID-19 n = 69 and pneumonia n = 9.

## Discussion

In the current study, a genome-based microbiome signature, which was composed of 33 HQMAGs at time of admission, was found to be associated with the severity and prognosis of COVID-19 in hospitalized patients. With these 33 genomes as reference, we were also able to validate the microbiome signature in datasets collected from three independent studies.

We came to this finding by way of a unique analytical strategy on the microbiome dataset. Previous studies relied on reference database to profile gut microbial composition at taxonomic levels and explored the relationships between different taxa and COVID-19^3–11^. Our strategy took a reference-free, discovery approach which does not need any prior knowledge. This allowed us to keep the novel part of the dataset intact. In addition, the use of high-quality draft genomes in our study ensured the highest possible resolution for identifying microbiome signature associated with COVID-19, which overcame the pitfalls of taxon-based analysis^14^. In the previous studies based on taxa-level, *Enterococcus faecium, Enterococcus* avium and *Akkermansia muciniphila* have been reported to be enriched in severe/critical COVID-19 patients and positively correlated with symptom seveity^3,9^. In our results, totally 28 A. *muciniphila, 2 E. avium and 5 E. faecium* HQMAGs were assembled in our dataset, but only 3 *E. faecium* and 1 each from *A. muciniphila* and *E. avium* enriched in severe/critical group, suggesting that not all strains from the 3 species were associated with COVID-19 severity. Another example is that *Faecalibacterium prausnitzii,* a key producer of SCFAs, is consistently depleted in COVID-19 patients and negatively correlated with disease severity^3,4^, but in our results, only half of the *F. prausnitzii* HQMAGs in our dataset were negatively associated with COVID-19 symptom severity. These indicate that the associations between gut microbiota and COVID-19 are strain/genome-specific. This means that even species does not have the necessary resolution to reveal associations of gut microbiome with COVID-19.

In our study cohort, RF classification models using the abundance of the 33 HQMAGs as input variables were able to discriminate the different symptom severity groups of COVID-19 patients at admission. Such a capacity of the 33 HQMAGs for discriminating COVID-19 symptom severities and distinguishing COVID-19 subjects from healthy subjects, non-COVID-19 and pneumonia controls was further validated in independent Chinese cohorts^7,9,44^. These indicate the feasibility to use this microbiome signature risk stratification of COVID-19 patients at least for Chinese cohorts. It will be worth validating the applicability of the microbiome signature in COVID-19 diagnosis in cohorts across ethnicity and geography.

In addition to identify COVID-19 associated gut microbiota at genome level, we used guild-based analysis to reveal potential interactions among key gut bacteria via co-abundance network. We found that genomes enriched in the mild/moderate group and genomes enriched in severe/critical group formed as two guilds, Guild 1 and Guild 2, respectively. The genomes in Guild 1 had higher SCFA producing genetic capacity while Guild 2 had more VF and ARG encoding genes. Reduced abundance of SCFA producing pathways has been correlated with more adverse clinical outcomes in COVID-19 patients^7^. The expression levels of VF and ARG, as measured by metatranscriptomic sequencing, were significantly higher in COVID-19 patients as compared with healthy controls and non-COVID-19 pneumonia controls^45^. Higher abundance of Guild 1 and lower abundance of Guild 2 was associated with reduced severity of our COVID-19 patients. Such a two competing guilds structure, in which one beneficial guild and one detrimental guild competed with each other and influenced host health, has been reported as a core microbiome signature associated with various chronic diseases^46^. The findings here suggest that such two competing guilds microbiome signature may be also applicable to infectious diseases.

The microbiome signature found in the current study was also associated with COVID-19 prognosis. The two competing guilds microbiome signature at admission was associated with several immune, coagulation, hemogram and biochemical indicators of hospitalized patients after 1 week. These indicators included D-Dimer, Fg, FDP, LYMPH%, LYMPH#, Neu%, TC, CA, ALB, TBIL, DBIL, LDH and glucose, which have been reported to play the essential role in the host immune response to COVID-19 infection and disease progression^25–41^. The microbiome signature may server as an early predictor of COVID-19 prognosis as it was positively associated with bio-clinical parameters that have inverse relationship with poor prognosis, and negatively associated with those that have direct relationship with poor prognosis. More importantly, early time point variations of the two competing guilds microbiome signature were correlated with the later changes of these prognosis related bio-clinical parameters. These results suggest that the dysbiosis of gut microbiota may play an essential role in trigging severe symptoms after patients are infected by SARS-CoV-2.

Early identification and treatment of high-risk patients is critical for improving prognosis of COVID-19 when the end of the pandemic is not in sight due to emerging of SARS-CoV-2 variants such as Omicron. At time of admission, screening hospitalized COVID-19 patients with our genome-based guild-level microbiome signature may facilitate early identification of those patients with high risk of more severe outcomes and put them under intensive surveillance and preventive care.

## Methods

### Ethics statement

This study procedure was reviewed and approved by the Ethics Committee of Shanghai Public Health Clinical Center ((SHPHC, NO. YJ-2020-S080-02), and informed written consents were signed from all subjects according to with the Declaration of Helsinki. All experimental procedures were performed in strict accordance with the guidelines for biosafety operation of SARS-CoV-2 Laboratories of the National Health and Family Planning Commission (No. 2020 [70]) and the Shanghai Municipal Health and Family Planning Commission (No. 2020 [8]). Table S1 sample collection and severity information.

### Subject recruitment and sample collection

This study was retrospectively conducted in Shanghai Public Health Clinical Center, a designated hospital for COVID-19 treatment in East China. In total of 337 COVID-19 patients were recruited in this study, all patients were typed and grouped based on clinical symptoms by senior clinicians in strict accordance with the criteria following Diagnosis and Treatment Plan for SARS-CoV-2 (Trial Version 7) issued by the General Office of the National Health Commission. The clinical data of the study subjects, including patient epidemiology (age, gender, disease classification, length of hospital stay, duration of disease, clinical outcomes), and respective clinical laboratory test results (hematologic, clinical chemistry, coagulation, immune inflammatory indices, and radiographic indications) were stored in a computerized database in the hospital medical record system. Stool sampling was collected within 48 hours after admission in all patients from May to September 2020, ensuring that all patients did not receive intervention from antiviral, antibiotic, probiotic, hormone and other drug interventions. About 100 mg of patient’s feces were collected in a stool collection tube and frozen immediately in a −80 °C freezer until processing.

### Clinical laboratory examination and data collection

All laboratory tests were conducted in the department of laboratory medicine in the Shanghai Public Health Clinical Center. Sysmex XN-1000 automated hematology analyzer (Hisense Meikang medical electronics, Shanghai Co., Ltd) and its supporting test reagents were used to analyze blood routine tests including white blood cell count (WBC), lymphocyte count (LPC), platelet count (PLT), Neutrophil (%), monocyte (%), lymphocyte (%), hemoglobin (HGB), Hypersensitive C-reactive protein (hs-CRP), etc. Biochemical parameters such as albumin (ALB), Amylase (AMY), cholinesterase (CHE), lactate (LACT), lactate dehydrogenase (LDH) alkaline phosphatase (ALP), glucose (GLU), creatinine (CRE), uric acid (UA), prealbumin (PA) were measured by a biochemical immunoassay workstation (ARCHITECT 3600J, Abbott Laboratories Co., USA). Urine routine (pH value, specific gravity, urobilinogen, leukocyte esterase, nitrite, urine protein, glucose, ketone body, bilirubin and occult blood) was measured by Cobas6500 urine dry chemical analysis system and supporting test strips (Roche, Switzerland). For the Coagulation indicators, The STA Compact Max was used to measure fibrinogen, DD-dimer, fibrinogen degradation products (FDP), prothrombin time (PT), activated partial thromboplastin time (APTT), thrombin time (TT), etc.

### Plasma cytokine measurements

FACS Canto II Flow cytometer (BD Biosciences, USA) was employed to the Lymphocyte analysis, CD3+ cell counts, CD4+ cell counts, CD8+ cell counts, CD19+ cell counts, CD16+ CD56+ cell counts, and CD4+/CD8+ percentage were detected. Plasma cytokine-related parameters including IL-1β, IL-2, IL-4, IL-5, IL-6, 1L-8, IL-10, IL-12P70, IL-17, tumor necrosis factor alpha (TNF-α), interferon-alpha (IFN-α), and interferon-gamma (INF-γ) were measured using a microsphere array kit by FACS Canto II cytometer (Raisecare Biotechnology, China).

### Gut microbiome analysis

#### DNA extraction and metagenomic sequencing

The laboratory procedure in this section was performed by trained laboratory personnel under the condition of tertiary protection in the bio-safety level laboratory-2 (BSL-2) qualified laboratory. The DNA was extracted from fecal samples using the bead-beating method as previously described ^50^, QIAamp PowerFecal Pro DNA Kit (Germany, QIAGEN) was employed to perform DNA extraction according to manufacturer’s instructions. Briefly, fecal sample (~ 100 mg) were dissolved by Powerlyzer lysate in PowerBead Pro Tube, vigorous shaking for 10 minutes, centrifugation. Total genomic DNA was captured on a silica membrane in spin-column. DNA is then washed and eluted, The A260/A280 ratio near 1.8, concentration and curve observations were used to identify DNA extraction quality assessment. Qualified DNA sample ready for downstream applications. Metagenomic sequencing was performed using Illumina Hiseq 3000 at GENEWIZ Co. (Beijing, China). Cluster generation, template hybridization, isothermal amplification, linearization, and blocking denaturing and hybridization of the sequencing primers were performed according to the workflow specified by the service provider. Libraries were constructed with an insert size of approximately 500 bp followed by high-throughput sequencing to obtain paired-end reads with 150 bp in the forward and reverse directions. Table S2 shows the number of raw reads of each sample.

#### Data quality control

Trimmomatic^51^ was used to trim low quality bases from the 3⍰ end, remove low quality reads and remove reads < 60bp, with parameters: LEADING:6 TRAILING:6 SLIDINGWINDOW:4:20 MINLEN:60. Reads that could be aligned to the human genome (H. sapiens, UCSC hg19) were removed (aligned with Bowtie2^52^ using --reorder--no-hd --no-contain --dovetail). Table S2 shows the number of high-quality reads of each sample for further analysis.

#### De novo assembly, abundance calculation and taxonomic assignment of genomes

De novo assembly was performed for each sample by using MEGAHIT^53^ (--min-contig-len 500, --presets meta-large). The assembled contigs were further binned using MetaBAT 2^54^ and MaxBin 2^55^. A refinement step was then performed using the bin_refinement module from MetaWRAP^56^ to combine and improve the results generated by the 2 binners. The quality of the bins was assessed using CheckM^57^. Bins had completeness > 95%, contamination < 5% and strain heterogeneity = 0 were retained as high-quality draft genomes (Table S3). The assembled high-quality draft genomes were further dereplicated by using dRep^58^. DiTASiC^59^, which applied kallisto for pseudo-alignment^60^ and a generalized linear model for resolving shared reads among genomes, was used to calculate the abundance of the genomes in each sample, estimated counts with P-value > 0.05 were removed, and all samples were downsized to 30 million reads (One sample at admission with read mapping ratio ~32%, which could not be well represented by the high quality genomes, were removed in further analysis). Taxonomic assignment of the genomes was performed by using GTDB-Tk^61^ (Table S4).

#### Gut microbiome functional analysis

Prokka^62^ was used to annotate the genomes. KEGG Orthologue (KO) IDs were assigned to the predicted protein sequences in each genome by HMMSEARCH against KOfam using KofamKOALA^63^. Antibiotic resistance genes were predicted using ResFinder^64^ with default parameters. The identification of virulence factors were based on the core set of Virulence Factors of Pathogenic Bacteria Database (VFDB^65^, download July 2020). The predicted proteins sequences were aligned to the reference sequence in VFDB using BLASTP (best hist with E-value < 1e-5, identity > 80% and query coverage > 70%). Genes encoding formate-tetrahydrofolate ligase, propionyl-CoA:succinate-CoA transferase, propionate CoA-transferase, 4Hbt, AtoA, AtoD, Buk and But were identified as described previously^66^.

#### Gut microbiome co-abundance network construction and analysis

Fastspar^67^, a rapid and scalable correlation estimation tool for microbiome study, was used to calculate the correlations between the genomes with 1,000 permutations at each time point based on the abundances of the genomes across the patients and the correlations with BH adjusted P < 0.05 were retained for further analysis. The co-abundance network was visualized using Cystoscape v3.8.1^68^

#### Definition of guild-level microbiome index (GMI)

We define the GMI using the abundance of the 33 MAGs and their relationships. For each individual samples, the GMI of sample j that was donated by GMIj was calculated as below:

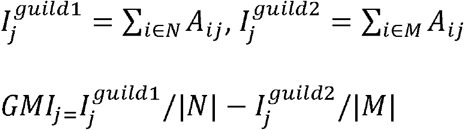

Where *A_ij_* is the relative abundance of HQMAG i in sample j. N and M are subsets of HQMAGs in guild 1 and guild 2 respectively. | N | and | M | are the sizes of these two sets.

#### Validation in an independent cohort

The metagenomic sequencing data from 24 mild/moderate and 14 severe/critical COVID-19 patient from the study was download from ENA database under PRJNA792726 (Table S5). The metagenomic sequencing data from 66 COVID-19 patients (first sample after admission), 69 age- and sex-matched non-COVID-19 controls and 9 community-acquired pneumonia controls but were negative for COVID-19 from the study conducted by Zhang et al.^7^ were download from ENA database under PRJNA689961 (Table S5). The metagenomic sequencing data from 46 COVID-19 patients and 19 healthy controls was download from ENA database under PRJEB4355 (Table S5). KneadData (https://huttenhower.sph.harvard.edu/kneaddata/) was applied to perform quality control of the raw reads with parameters: --decontaminate-pairs strict, --run-trim-repetitive, --bypass-trf, --trimmomatic-options= “SLIDINGWINDOW:4:20 MINLEN:60”. Reads that could be aligned to the human genome were identified and removed in KneadData by aligning reads against Homo sapiens hg37 genome. The abundance of the 33 MAGs were estimated by using Coverm v0.6.1 (https://github.com/wwood/CoverM) with parameters: coverm genome --min-read-aligned-percent 90 --min-read-percent-identity 99 -m relative_abundance.

### Statistical Analysis

Statistical analysis was performed in the R environment (R version4.1.1). Kruskal-Wallis test followed by Dunn’s post hoc (two-sided) was applied to compare the different severity groups. Redundancy analysis was conducted based on the Hellinger transformed abundance to find Specific gut microbial members associated with COVID-19 severity. A marginal PERMANOVA test including both age and symptom severity was used to compare the overall gut microbial composition. Random Forest with leave-one-out cross-validation was used to perform regression and classification analysis based on the microbiome signature and clinical parameters/groups.

## Acknowledgements

This work was supported by grants from the National Natural Science Foundation of China (No. 8210061470), the Shanghai Sailing Program (No. 21YF1438800) and Notitia Biotechnologies Company. The computations in this paper for the discovery cohort were run on the *π* 2.0 cluster supported by the Center for High performance Computing at Shanghai Jiao Tong University. The authors acknowledged the Office of Advanced Research Computing (ORAC) at Rutgers, The State University of New Jersey for providing access to the Amarel cluster and associated research computing resources that have contributed to the computations in this paper for validation cohorts.

## Author contributions

L. Z., C.Z., and Z.Z. designed and supervised the study. M.G. and G.W. performed the majority of experiments and conducted the analyses. Y.T., Y. L. and X.G. participated in the experiment guidance, X. J. and W.Q. performed and collected the laboratory detection. L.Z., C.Z., G.W. and M. G. wrote and revised manuscript. All authors contributed to the article and approved the submitted version. M.G. and G.W. contributed equally to this work.

## Competing financial interests

L.Z. is a co-founder of Notitia Biotechnologies Company.

## Data availability

The metagenomic sequencing data to the current study are freely available upon request to corresponding authors.

## Code availability

Parameters of the bioinformatic tools applied in the study were showed in the method section. Scripts and command lines related to the current study are freely available from the corresponding authors upon request.

## Ethics & Inclusion statement

We have carefully considered research contributions and authorship criteria when involved in multi-region collaborations involving local researchers so as to promote greater equity in research collaborations.

## Notes

### Competing Interest Statement

Liping Zhao is a co-founder of Notitia Biotechnologies Company.

