## Supplemental Figures for "Guild-level microbiome signature associated with COVID-19 severity and prognosis"

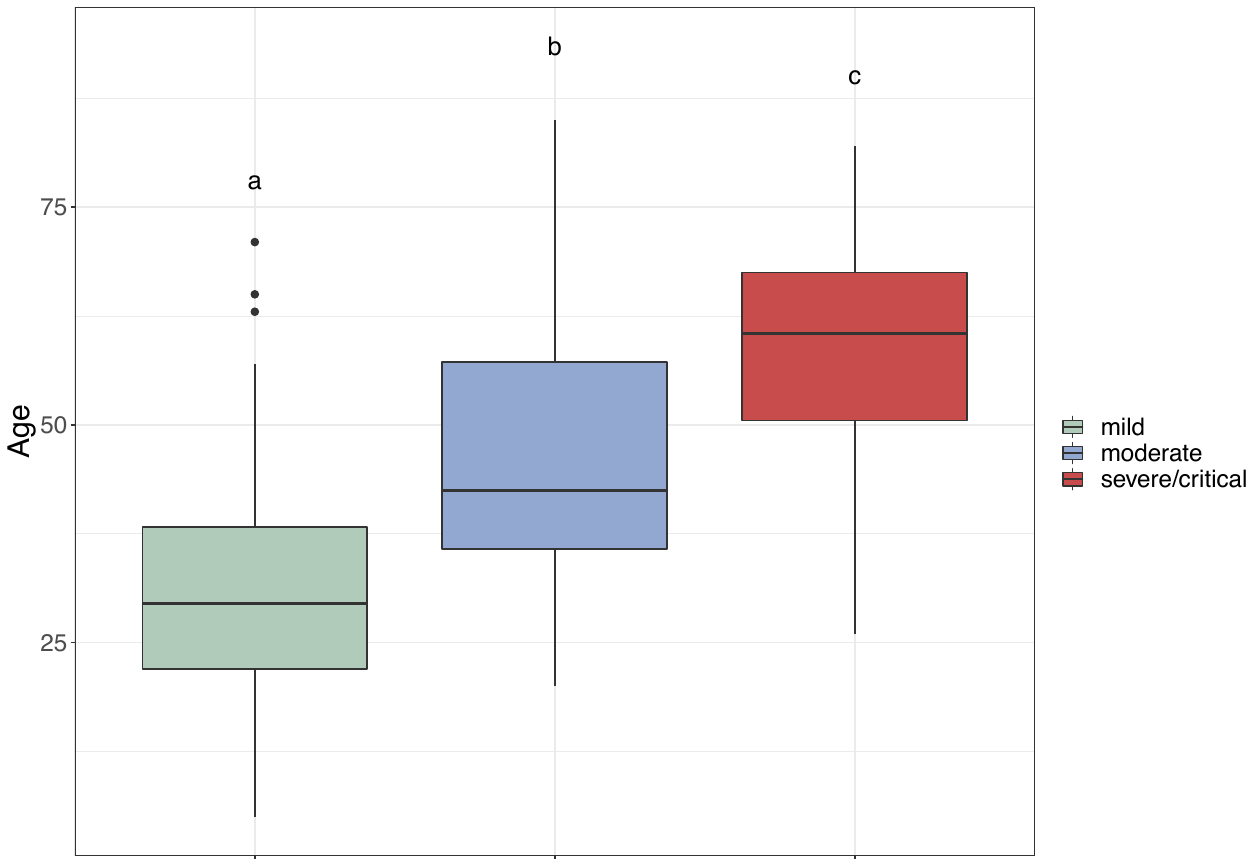

**Figure S1. COVID-19 Patient with more severe symptoms were older.** Data points not sharing common compact letters were significantly different from each other (p < 0.05). Boxes show the medians and the interquartile ranges (IQRs), the whiskers denote the lowest and highest values that were within 1.5 times the IQR from the first and third quartiles, and outliers are shown as individual points. Kruskal-Wallis test followed by Dunn’s post hoc (two-sided) was applied to compare the groups. Compact letters reflect the significance of the test (P < 0.05). Mild: n = 88; moderate n = 196, severe/critical n = 12.

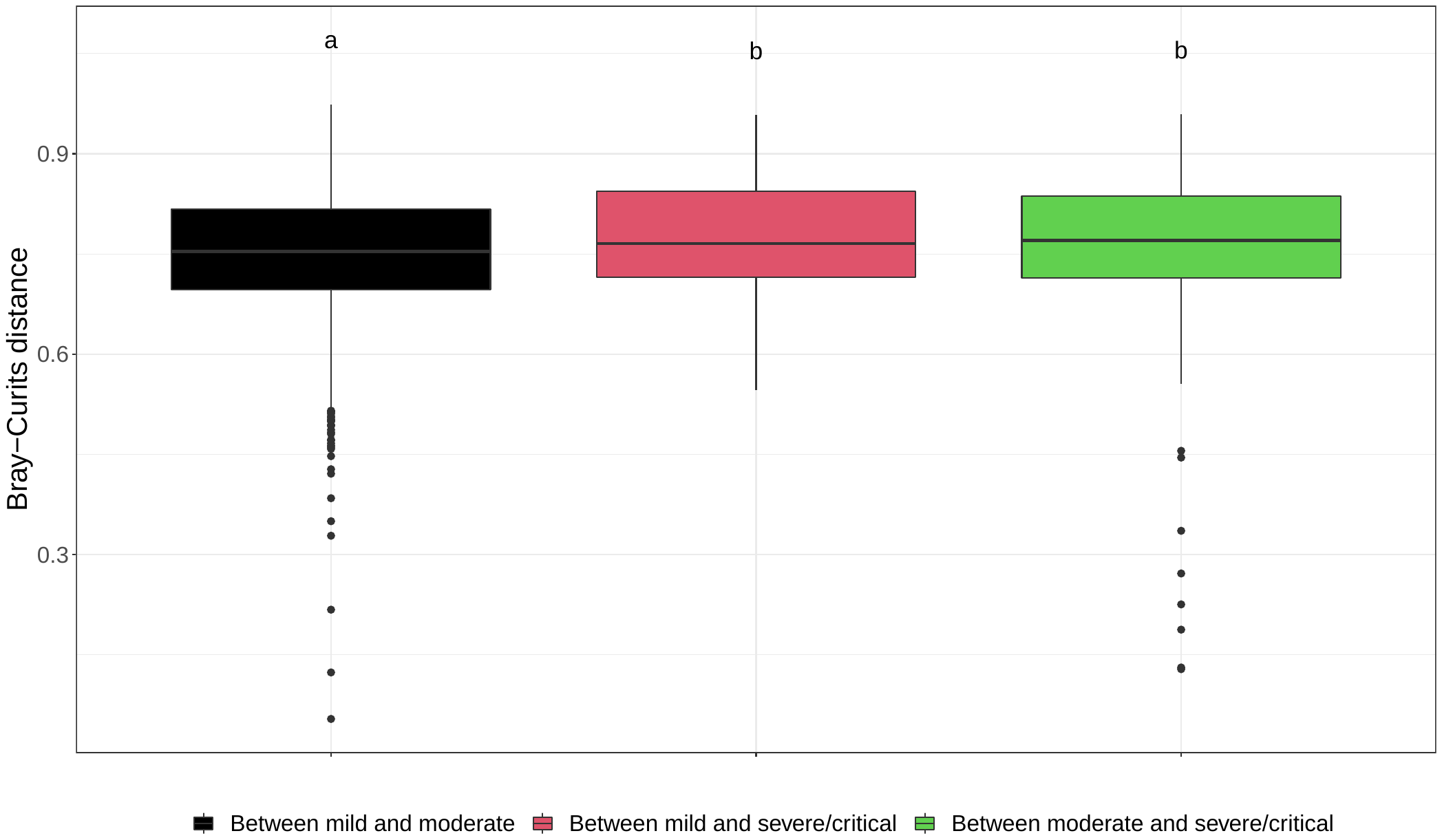

**Figure S2. Between-group Bray-Curtis distance.** Data points not sharing common compact letters were significantly different from each other (p < 0.05). Boxes show the medians and the interquartile ranges (IQRs), the whiskers denote the lowest and highest values that were within 1.5 times the IQR from the first and third quartiles, and outliers are shown as individual points. Kruskal-Wallis test followed by Dunn’s post hoc (two-sided) was applied to compare the groups. Compact letters reflect the significance of the test (P < 0.05).

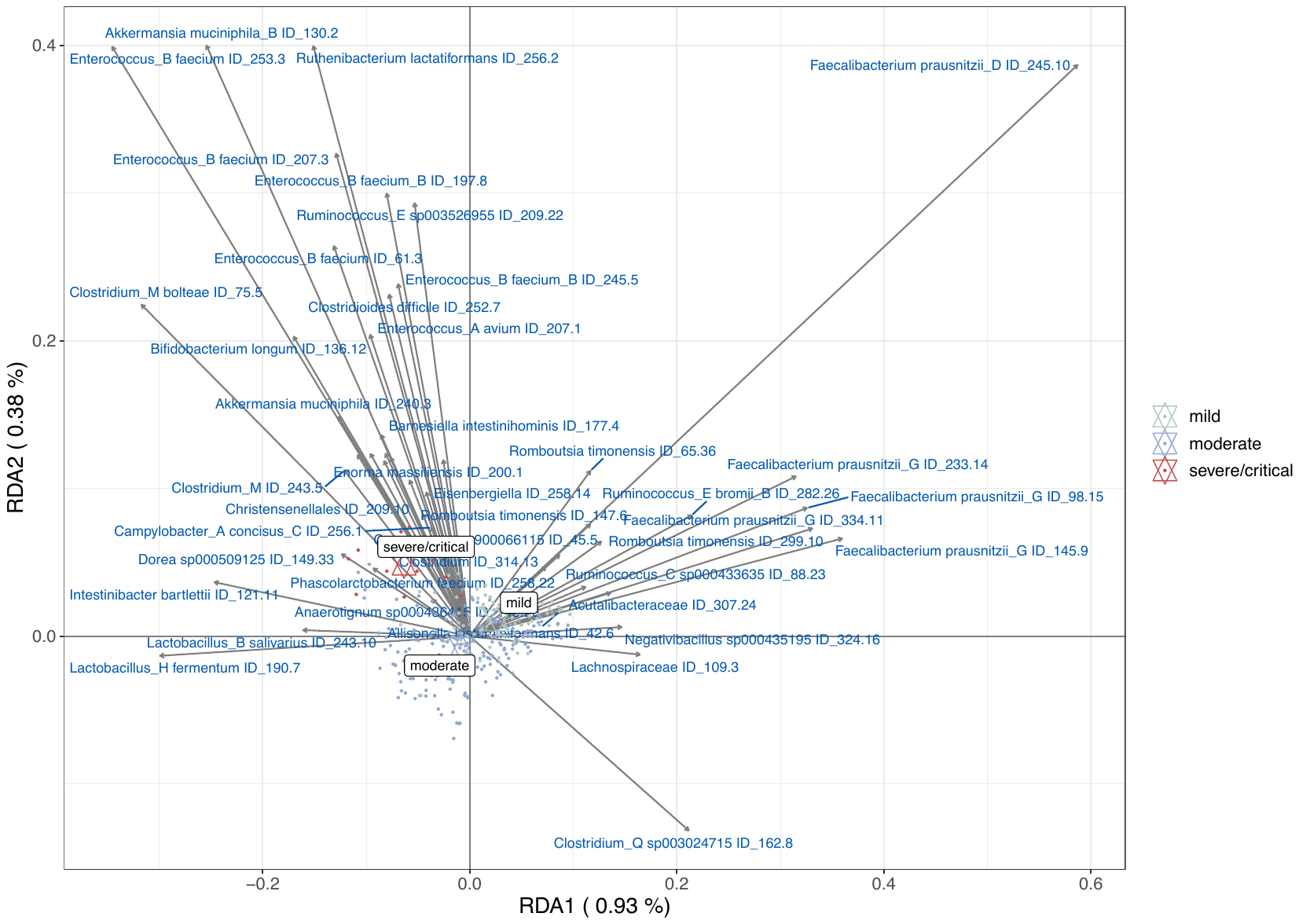

**Figure S3. Triplot of redundancy analysis (RDA) of the microbial composition based on the 2,568** HQMAGs**.** The three symptom severity groups were used as environmental variables. Samples are indicated by dots. HQMAGs with at least 5% of the variability in their abundance explained by RDA1 and RDA2 are indicated by blue arrows. RDA analysis was conducted based on the Hellinger transformed abundance.

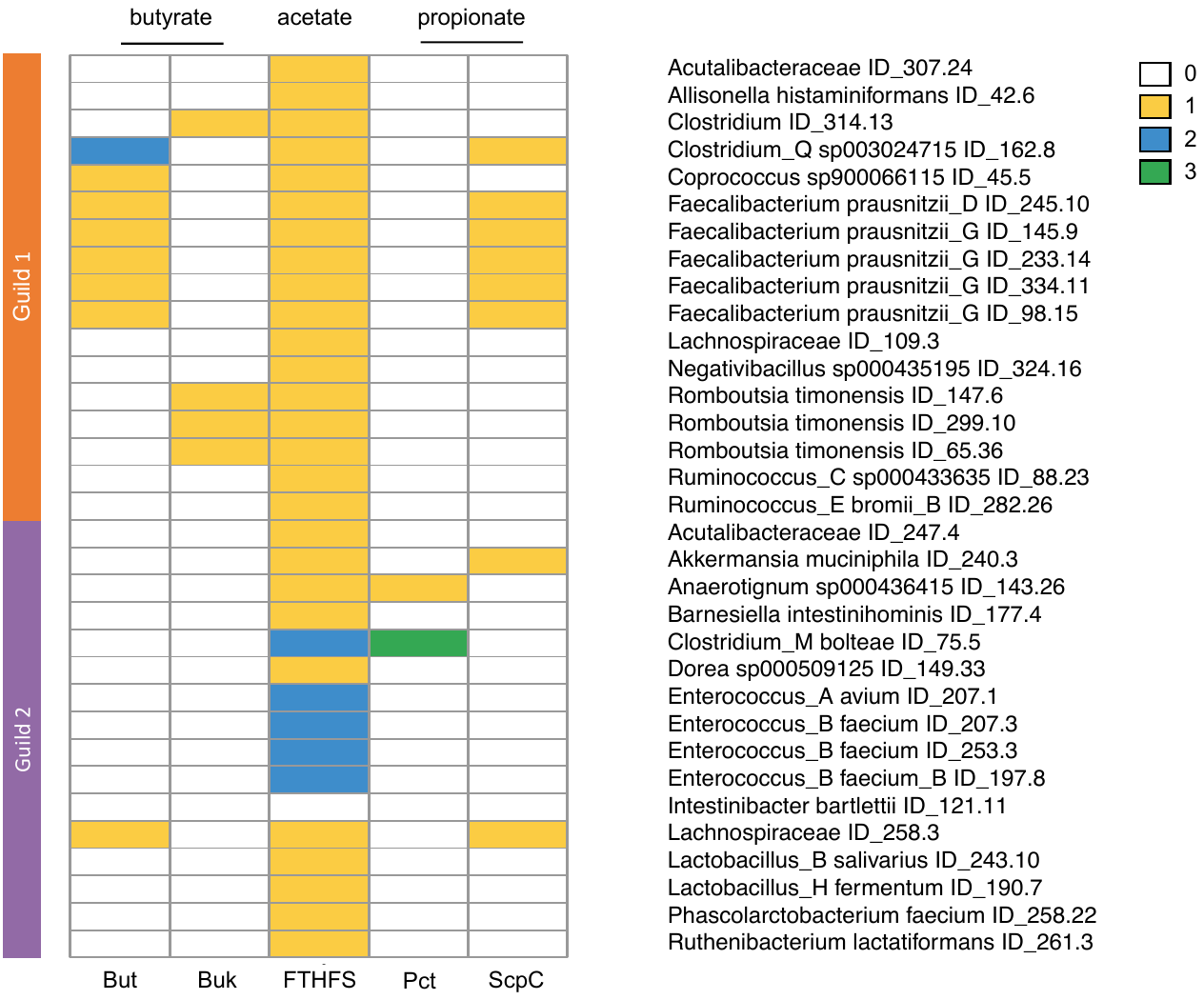

**Figure S4. Differences in genetic capacity of short chain fatty acid biosynthesis between the 2 guilds.** The heatmap shows the gene copy number of But: Butyryl–coenzyme A (butyryl-CoA): acetate CoA transferase, Buk: butyrate kinase; FTHFS: formate-tetrahydrofolate ligase for acetate production; ScpC: propionyl-CoA succinate-CoA transferase and Pct: propionate-CoA transferase for propionate production

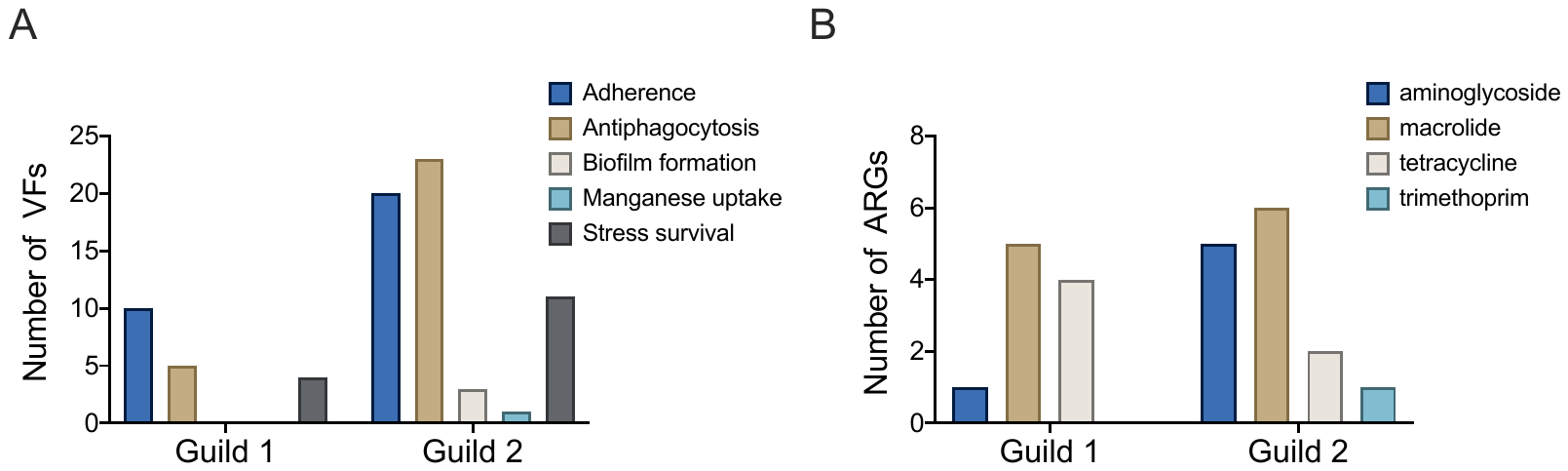

**Figure S5. Differences in genetic capacity for pathogenicity and antibiotic resistance between the 2 guilds.** (A) The bar plot shows the number of genes encoding virulence factors (VF) and classes of VFs. (B) The bar plot shows the number of ARGs and the corresponding antibiotic resistance types.

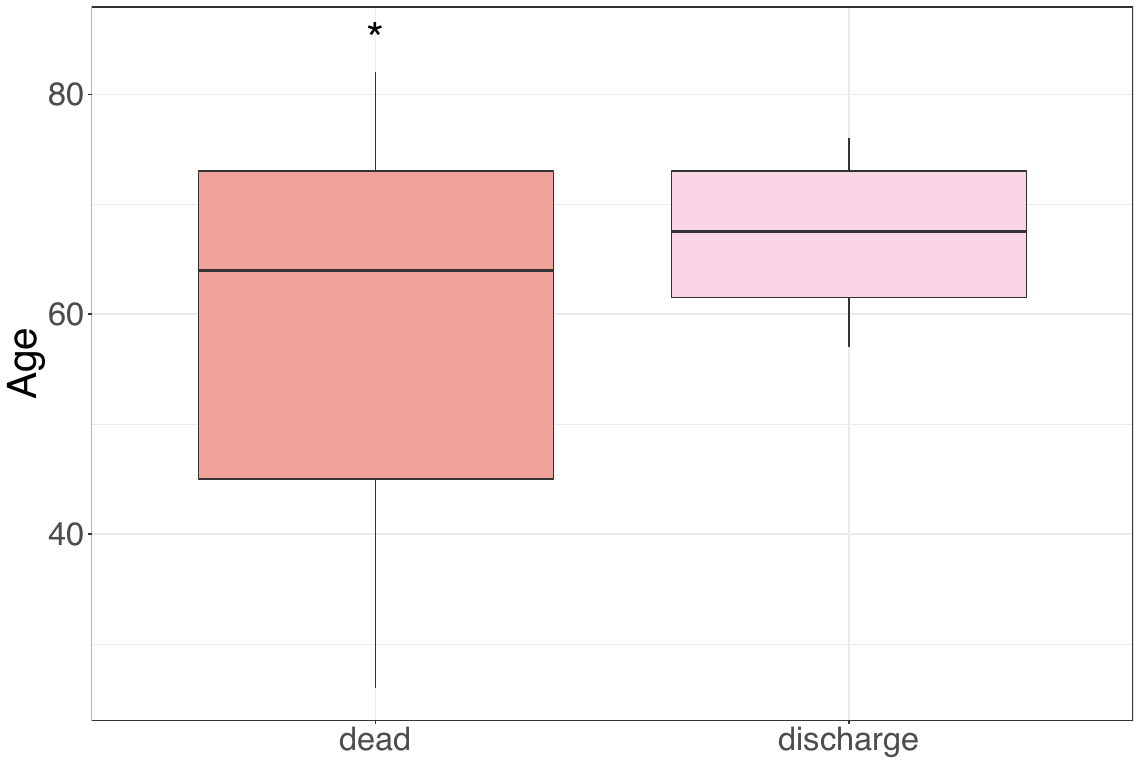

**Figure S6. In critical COVID-19 patients at admission, the age of dead patients was significantly smaller than those were discharged**. Mann-Whitney test (two-sided) was used. N = 3 for dead, N =4 for discharge. * P < 0.05

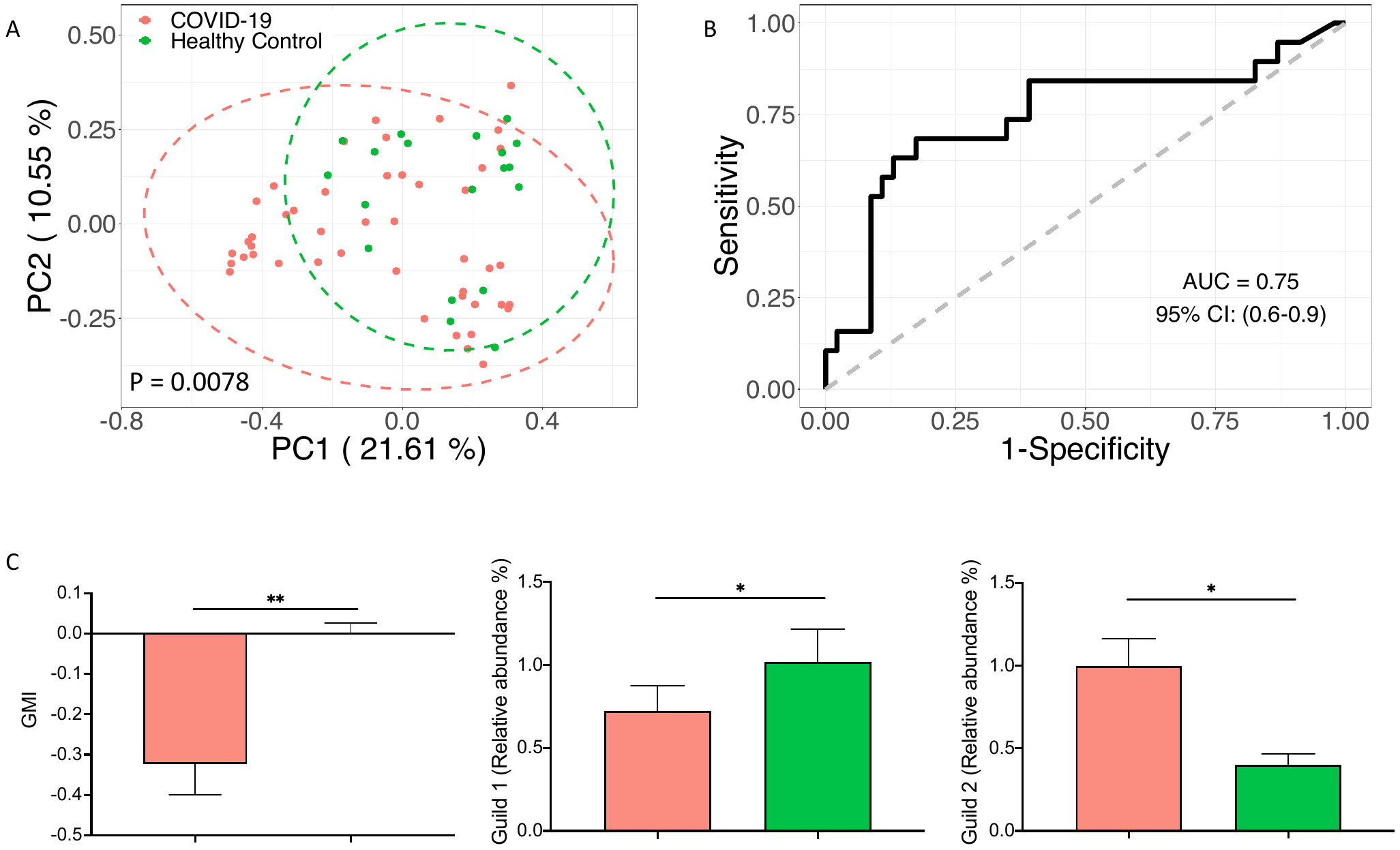

**Figure S7. The genome-based microbiome signature enables to classify COVID-19 from heathy control in an independent dataset.** (A) Principal Coordinate Analysis based on Bray-Curtis distance calculated from the abundance of the 33 MAGs. PERMANOVA test showed significant differences in the composition of the 33 MAGs between the two groups(B) The area under the ROC curve (AUC) of the Random Forest classifier based on the 33 MAGs to classify COVID-19 and healthy subjects. Leave-one-out cross validation was applied. (C) Significant differences in Guild-level Microbiome index (GMI) and abundances of Guild 1 and Guild 2 between COVID-19 and healthy subjects. The barplot summarized the mean and S.E.M. Mann-Whitney test (two-sided) was applied to compare the groups. COVID-19 n = 46, Healthy Control =19. * P < 0.05, ** P < 0.01

**Table S1 Subject grouping information**

[In the Excel file]

**Table S2 Raw and high-quality reads of each sample.**

[In the Excel file]

**Table S3 The genome quality assessed by CheckM.**

[In the Excel file]

**Table S4 The taxonomic assignment of genomes by GT-DBTK.**

[In the Excel file]

**Table S5 The information of samples for validation in the independent datasets.**

[In the Excel file]
